# Virtual manipulation of bird tail postures demonstrates drag minimisation when gliding

**DOI:** 10.1101/2020.06.09.141994

**Authors:** Jialei Song, Jorn A. Cheney, James R. Usherwood, Richard J. Bomphrey

**Affiliations:** School of Mechanical Engineering, Dongguan University of Technology, Dongguan, Guangdong, China; Structure and Motion Laboratory, Royal Veterinary College, North Mymms, Hatfield, Hertfordshire, United Kingdom

**Keywords:** tail function, drag reduction, efficiency, bird flight

## Abstract

Aerodynamic function of the avian tail has previously been studied using observations of bird flight, physical models in wind tunnels, theoretical modelling and flow visualization. However, none of these approaches has provided rigorously quantitative evidence concerning tail functions since: appropriate manipulation and controls cannot be achieved using live animals; and the aerodynamic interplay with wings and body challenges reductive theoretical or physical modelling approaches. Here, we have developed a comprehensive analytical drag model, calibrated by high-fidelity computational fluid dynamics (CFD), and used it to investigate the aerodynamic action of the tail by virtually manipulating its posture. The bird geometry used for CFD was reconstructed previously using stereo photogrammetry of a freely gliding barn owl and validated against wake measurements. Using this CFD-calibrated drag model, we predicted the drag production for 16 gliding flights with a range of tail postures. These observed postures are set in the context of a wider parameter sweep of theoretical postures, where the tail spread and elevation angles were manipulated virtually. The observed postures of our gliding bird corresponded to near minimal total drag.

**Author Summary:** The aerodynamic contribution of bird tails is challenging to study; strong interactions between wings, body and tail make models isolating the contributions of different body parts difficult to interpret. Further, methods for direct manipulation are limited, and confounding compensation is likely in live, free-flying birds. To circumvent these issues, we applied high-fidelity CFD to a range of measured gliding owl geometries in order to develop a comprehensive analytical drag model. This enabled the drag implications of virtually-manipulated tail postures to be explored. The theoretical postures that cause minimum drag match those used by owls. The drag model demonstrates the importance of the viscous component of drag, which is of particular relevance to fliers at the scale of birds and, increasingly, smaller UAVs.

## Introduction

Avian tails may have many aerodynamic functions during steady gliding. It has been demonstrated that the tails can be used to maintain stability and balance as well as control turning [1-3]. In slow flight, tails spread and pitch downward producing lift, which supplements that from the wings to support body weight, and can also stabilize the pitching moment [4-6]. In sideslip, tails provide a yaw moment, making a significant contribution to yaw stability [7]. Tails can also be configured to reduce drag and increase glide efficiency. Classical wing theory indicates that, for wings of finite span and no interruption from a fuselage or body, the ideal shape to minimise induced drag is an elliptical wing with constant downwash across the span [8]. However, the torso does interrupt the wings and disrupts the gradual change in wing profiles along the span, likely causing a deficit of downwash in the central region behind the body. A tail could potentially compensate for diminished downwash in this region by smoothing or equalising spanwise variation in the downwash and thus reducing the inviscid, induced, drag component of the total drag burden [9]. If the tail is held at a higher angle of attack than that which would equalise the downwash, the higher lift can act to reduce total drag further by reducing the viscous drag component [10]. This is particularly important at the small sizes and slow speeds of birds and runs contrary to the idea that tails are used to minimise the inviscid (or induced) component of drag alone. Tails also alter the flow over the dorsal body surface and inhibit flow separation behind the body, reducing the drag of the body [3,11].

Due to strong interactions between birds’ wings, torso and tail, and the difficulties of controlling posture in flight experimentally, direct study of the aerodynamic functions of avian tails is challenging using cursory theoretical models or quantitative experimental measurement. High-fidelity computational fluid dynamics (CFD) modelling provides a new approach for studying the aerodynamic functions of tails because it can determine the detailed aerodynamic implications of a variety of virtually manipulated tail postures. By testing observed configurations adopted by birds and also sweeping through a range of virtual configurations that we do not observe, we can contextualise the natural geometries and identify where they appear on an optimality landscape.

### Overview

Using CFD to understand the three-dimensional aerodynamics of birds is computationally burdensome. High-fidelity CFD still requires significant time investment for each condition. This issue is compounded by our knowledge that tail manipulations affect both lift and drag and, in order to make meaningful comparisons between configurations, an iterative approach is necessary to maintain lift that balances body weight. Comparing configurations that support equal lift is particularly critical when the question of interest is related to drag minimization. For these reasons, we develop a model to reduce the total number of simulations and interpolate the lift and drag, with all unknown parameters obtained directly or fitted from CFD calculations for each geometry.

In this study, we describe a new drag model and parameterise it with high-fidelity CFD simulations. We solved the Navier-Stokes equations for fluid motion around the detailed geometry of birds acquired using multi-camera photogrammetry. These high-fidelity simulations were previously validated by large volume quantitative wake visualisations [12]. With the drag model, we calculated the best aerodynamic performance (defined as minimum drag) of the entire bird with a range of different tail postures (varying spread and pitch) while producing constant lift. We then compared the predicted tail posture to our observations of 16 gliding flights performed by a barn owl within a large laboratory facility.

### Drag model

Drag is a critical parameter to understand because it has a direct effect on flight performance and, consequently, a bird’s ecological limits. One common – perhaps prevailing - drag model used for bird flight estimates [13] is derived from standard aeronautical practice. It comprises three elements: induced drag (D_ind_), profile drag (D_pro_) and parasite drag (D_par_), which are summed:

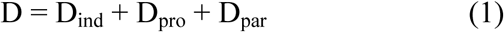

Induced drag arises when the wings and tail produce the lift required to support the bird’s weight, which behaves as:

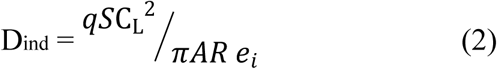

where 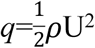 is dynamics pressure, C_L_ is the lift coefficient, *AR = b*_*2*_ */S* is the wing aspect ratio (b is wing span and *S* is planform area), e_i_ is the span efficiency (describing the increase in drag due to a departure from the ideal, even, downwash distribution across the wing span, Eq. 10 in Materials and Methods). Profile drag is due to skin friction and the pressure difference between the downstream and upstream surfaces, written as

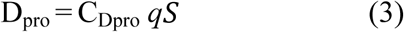

where C_Dpro_ varies somewhat with lift [14,15], but has often been approximated as constant of 0.02 [16-18]. Parasite drag is associated with drag of the body, which behaves as

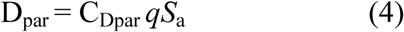

where *S*_a_ is frontal area and C_Dpar_ is a coefficient that is also sometimes treated as a constant value of 0.1 [5,19]. However, this also varies with factors such as airspeed and body angle [14].

Drag models that treat C_Dpro_ and C_Dpar_ as constants (Model I, Fig. 1) result in a gradual increase in drag with lift coefficient, with the increase due to the inviscid, induced drag component (Fig. 1). A recent implementation of the drag model [10] emphasised the quadratic rise in viscous, profile drag with lift (Model II, Fig. 1). In this model, total drag is again calculated using three additive terms: induced drag, viscous drag and a minimum drag coefficient which is assumed to occur at zero lift:

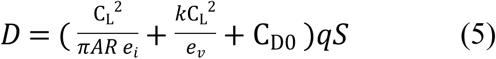

where the viscous efficiency coefficient:

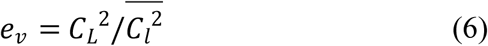

describes the increase in drag due to inefficient loading across the span: for a wing of uniform sectional shape this is a departure from a constant, sectional-lift-coefficient distribution across the span. For birds, especially those that are small and/or slow flying (and thus operating at low Reynolds numbers where viscous effects are more important), adopting a shape and pose that minimises viscous drag is likely to be close to the minimal total drag solution. This model explains the high tail angle of attack of three raptors (barn owl *Tyto alba*, tawny owl *Strix aluco* and goshawk *Accipiter gentilis*) and the consequent unexpectedly strong downwash beneath the tail [10].

**Figure 1.**
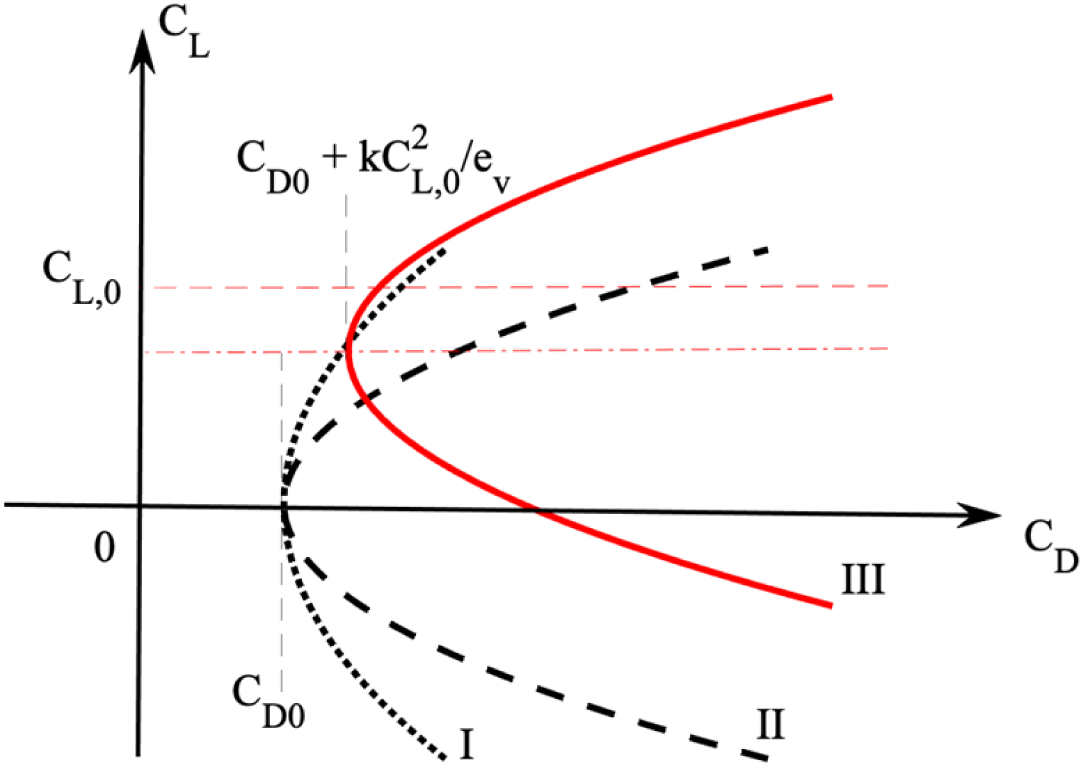
Sketch of different drag models. Model I (dotted line) assumes C_pro_ and C_par_ to be constant, with a dependency on lift entirely due to induced drag. Model II (dashed line) assumes no camber effect. Model III includes the camber effect.

Model II provides an approximation that is valid for wing sections with negligible camber– those for which the minimum drag occurs at minimum lift; however, avian wings are usually highly cambered [20-23]. To expand the model to cover cambered wings requires an additional parameter, C_L,0_. Camber offsets the minimum-drag condition to a point where positive lift is produced [24]. To determine this new camber-induced-lift parameter, we used CFD to derive the value and assign it to our expanded model (Model III, Fig.1). For each 2-dimensional chord on the wing, the drag and lift can be approximated by the relationship [25]:

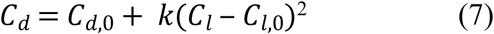

Where *C*_*l*_, *C*_*l*,0_ are the 2-dimensional lift coefficient and the lift coefficient at minimum drag, respectively, and *k* is a constant that defines the quadratic rise in *C*_*d*_ with *C*_*l*_ − *C*_*l*,0_. Then, using linear summation of induced drag and viscous drag, and the quasi 2-dimensional assumption, the overall drag of a 3-dimensional aerofoil behaves as:

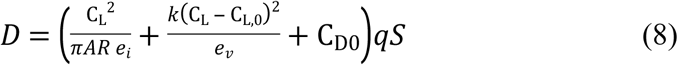

The first term within the brackets is the induced drag contribution, the second is the viscous drag, and the third is the minimum drag without camber effect.

### CFD modelling and its robustness

The high-fidelity bird surface mesh for CFD is based upon the geometry of a trained adult female barn owl, *Tyto alba*, weighing 340 g (Fig. 2). The owl’s geometry was obtained by cutting-edge stereo-photogrammetry [12], using a similar approach to that of Durston and colleges[26]. In the specific flight providing the baseline geometry for this CFD study, the bird flew at U = 7.88 m/s with a glide angle of *θ* = 2.9°, which is close to the average of all sequences and negligible acceleration in all axes [12]. Further bird model geometry information is shown in Table 1. The mesh vertices represent the point cloud closely, with 96% mesh vertices are within 3 mm of the point cloud and 90% are within 2 mm (Figure S1) (Video S1 shows the error distribution on bird surface). An additional 15 steady glides were measured for subsequent tail posture comparison with the minimal drag configuration predicted by our drag model.

**Table 1.** Parameters of the barn owl, *Tyto alba*.

| Parameters | values |
| --- | --- |
| Body mass, $m$ | 340 grams |
| Wing span, $b$ | 0.87 m |
| Planform area, $S$ | 0.1478 m<sup>2</sup> |
| Frontal area, $S_b$ | 0.0095 m<sup>2</sup> |
| Average chord length, $\bar{c}$ | 0.17 m |
| Aspect ratio, $AR$ | 5.12 |
| Flight speed, $U$ | 7.88 m/s |
| Gliding angle, $\theta$ | 2.9° |

**Table 3.** Mesh convergence study. The first layer thickness was changed while the inflation thickness was maintained. The use of *δ*_*t*_ = 0.1 mm gives 1.55% and 0.25% difference from the finest mesh (*δ*_*t*_ = 0.08 *mm*).

| First layer thickness | Lift | Drag |
| --- | --- | --- |
| $\delta_t = 0.08 \text{ mm } (y^+ \sim 2)$ | 3.22 | 0.2834 |
| $\delta_t = 0.1 \text{ mm } (y^+ \sim 3)$ | 3.17 (1.55%) | 0.2827 (0.25%) |
| $\delta_t = 0.2 \text{ mm } (y^+ \sim 7)$ | 3.15 (2.17%) | 0.2925 (3.21%) |

**Figure 2.**
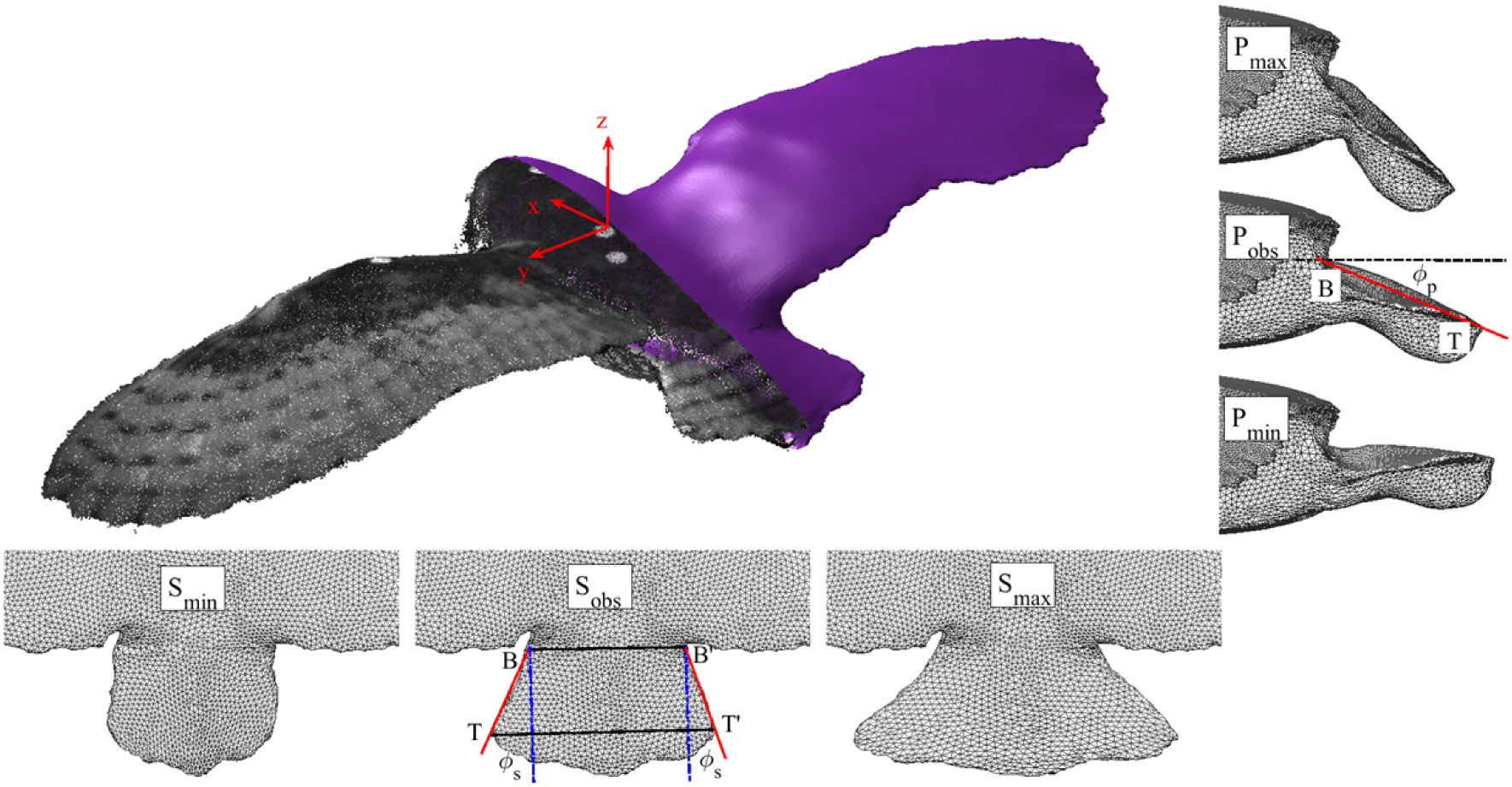
Point cloud obtained from stereo-photogrammetry and the surface mesh used for CFD simulations. The tail spread and pitch could be manipulated based on four landmarks. B and B’ determine the tail base and are defined as the points that give a minimal length when seen in vertical projection. Conversely, the tail tips T and T’, are defined by a connecting line that gives a maximal length when seen in vertical projection. The tail side edges are BT and B’T’. The projection of the angle between the side edges and glide trajectory on the sagittal plane gives the tail pitch angle, *ϕ*_*p*_. The projection of the side edges when viewed from above gives the tail spread angle, *ϕ*_*s*_. As the real tail is not perfectly symmetrical, we use the average of the two side edge angles to calculate both the spread and pitch angles. S_obs_ shows the observed tail spread angle (*ϕ*_*s*_ = 17.5°), while S_min_ and S_max_ shows the manipulated minimal and maximal spread. P_obs_ shows observed tail pitch angle (*ϕ*_*p*_ = 26.0°), while P_min_ and P_max_ shows the virtually manipulated minimal and maximal pitch.

Using the surface mesh reconstructed from the observed bird geometry, we virtually manipulate the tail posture’s spread angle, *ϕ*_*s*_, and pitch angle, *ϕ*_*p*_. The schematic in Fig. 2 shows the definition of these angles, highlighting the observed bird geometry and the limits of the manipulated postures in our parameter sweep. For the observed case, the tail spread angle was *ϕ*_*s*_ = 17.5° and the tail pitch angle was *ϕ*_*p*_ = 26.0°. We independently simulated tail spread angles from 0° < *ϕ*_*s*_ < 41° with six intervals and tail pitch angles from 6° < *ϕ*_*p*_ < 46° with seven intervals (see Video S2). These ranges comfortably envelop our observations of the gliding owl. We also simulated combinations of these parameters to test for interactions, giving a total of 42 tail postures.

The flow around the gliding barn owl was simulated using commercial CFD software FLUENT 19.1 (ANSYS, Inc., Canonsburg, PA, USA) that solves the incompressible viscous governing equations of the fluid. The flow regime for this bird is expected to be turbulent, with Reynolds number ∼88000 using the mean wing chord 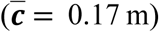 as our characteristic length. In order to simulate the flow accurately with a reasonable grid resolution, we used the *k* − *ω* SST turbulence model, which combines the advantages of the standard *k* − *ω* model and standard *k* − *ε* model, to give an excellent predictor of separated flows and adverse pressure gradient [27]. However, in some circumstances, results can potentially be sensitive to the choice of turbulence model so we compared the results from our *k* − *ω* SST simulations with several other popular models (Figure S2). The alternative turbulence models show a maximum of 6.25% lift and 15.6% drag differences compared with the *k* − *ω* SST model. While we can proceed with some degree of confidence that the turbulence models are in broad agreement with each other, it is important to be mindful that none of these turbulence models was specially designed for the Reynolds numbers or geometries of flying birds, and the cautious approach requires experimental validation of the CFD simulations. To achieve this, we measured the wake behind the same gliding barn owl using large-volume particle tracking velocimetry (PTV), which tracked over 20,000 neutrally buoyant soap bubbles and the detailed validation is described elsewhere [12]. Further detail of the photogrammetry and CFD setup are described in Materials and Methods.

## Results

### Force production and flight efficiency of virtually manipulated tail postures

The tail has many functions, one of which is to provide lift to supplement body weight support; this is especially important at low speeds [4-6, 28]. For the tail posture and flight speed we observed, the tail alone supports just 3% of body weight based on the integrated pressure distribution over the upper and lower surfaces of the tail feathers. However, by changing the tail’s spread and pitch, the lift generation of the entire bird varies dramatically, ranging from 84% (*ϕ*_*s*_ = 42°, *ϕ*_*p*_= 6°) to 122% (*ϕ*_*s*_ = 42°, *ϕ*_*p*_= 46°) of body weight (Fig 3a). Even though the lift produced by the tail alone is small, the tail alters the flow around the bird’s body and the proximal sections of the wings, conferring substantial control authority. The tail therefore acts as an aileron effectively changing the aerodynamics of the entire lifting surface (Video S3). With changes in tail spread and pitch, the total drag on the bird also changes (Fig. 3b). Drag increases with both spread and pitch of tail as *ϕ*_*p*_> 13°, and ranges from 88% to 172% of the observed case. When *ϕ*_*p*_< 13°, the tail tilts up, the resultant reversed camber running from beak to tail tips causes increased drag.

**Figure 3.**
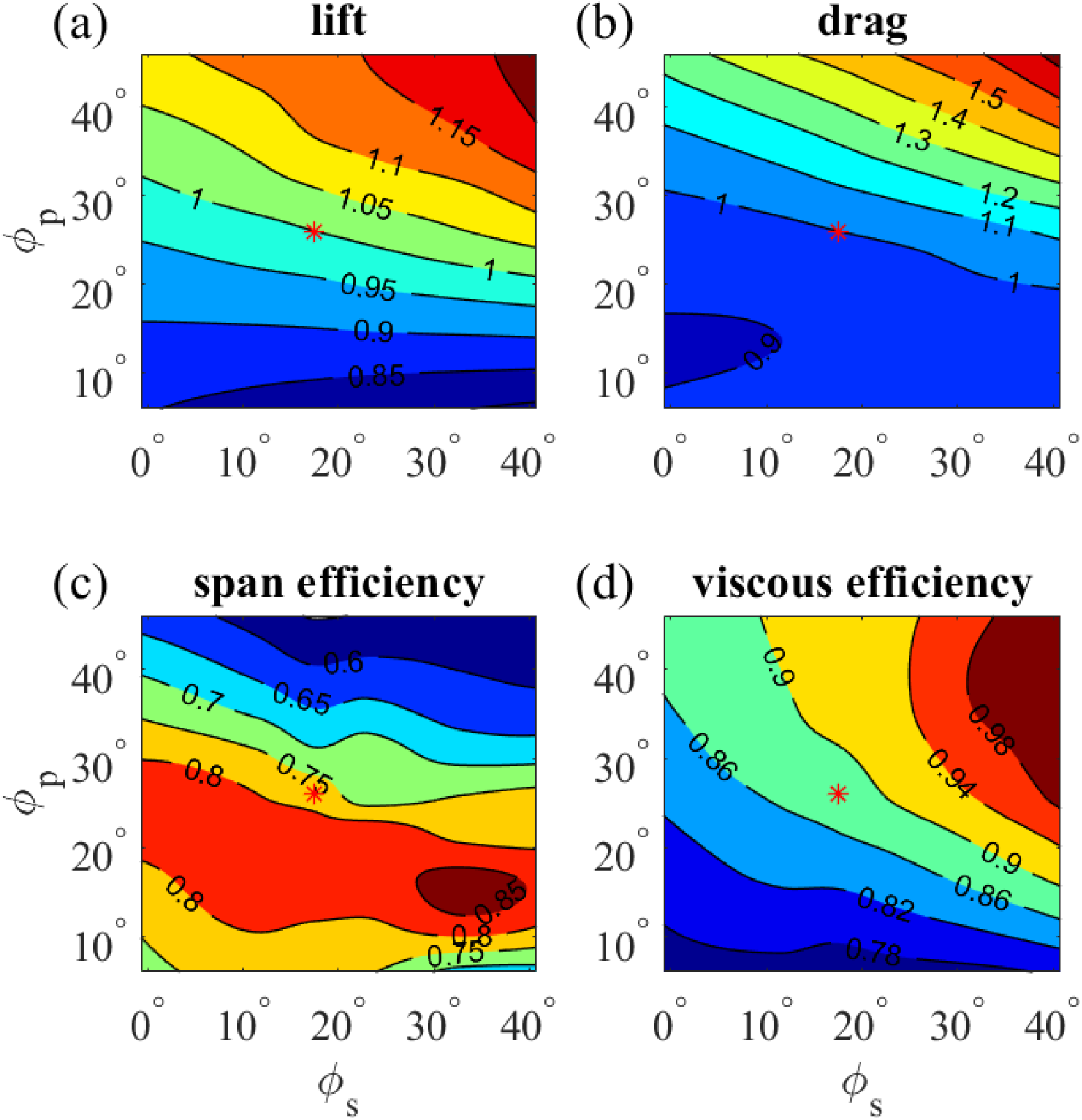
Forces and efficiencies across manipulations of tail spread and pitch. Normalized lift (a) and drag (b) relative to observed tail posture. Span (c) and viscous (d) efficiency.

As tail posture changes, the span loading and downwash distribution behind the bird varies. As is shown in Fig. 3c, within the range of tail spread and pitch, inviscid span efficiency, e_i_ ranges from 0.55 to 0.86, with the highest value obtained when the tail was widely spread with a moderate pitch angle (*ϕ*_*s*_= 36° and *ϕ*_*p*_ = 15°). The lowest span efficiency values occur when the tail is pitched down (depressed) to a large extent. The viscous efficiency factor is an indication of the evenness of the spanwise distribution of lift coefficient. Fig. 3d shows viscous efficiency factor with different tail postures, ranging from 0.78 to 0.99. Low e_v_ values are found when the tail is widely spread and pitched up (held high), whereas higher e_v_ values appear when the tail is widely spread and pitched-down. Importantly, the region of high e_v_ is not consistent with the region of high e_i,_ as is described in the introduction.

### Drag prediction with constant weight support

Steady gliding flight requires lift force to balance body weight. The components of drag change with tail spread and pitch but so does lift (Fig. 3). To compare like-for-like, we should compare peaks in drag efficiency when the bird is supporting body weight. Assuming flight speed remains constant, the whole-bird angle of attack can be altered in order to maintain constant lift; if we reduce angle of attack, we expect a concomitant reduction in drag.

To estimate the change in drag with constant weight support, we use drag model III, which includes the effect of camber (Eq. 8). In model III, the coefficient, *k*, is an unknown parameter that can only be obtained accurately from the C_L_-C_D_ polar plot by fitting a quadratic function. By changing the angle of attack of the entire bird (with its observed tail posture), we obtained five C_L_ *vs* C_D_ points (Fig. 4). Fitting these points with a binomial gives C_D_ = 0.206 C_L_^2^ – 0.113 C_L_ + 0.0526 for the three coefficients, respectively. Assuming e_i_ and e_v_ do not vary with angle of attack, the comparison between Eq. 8 and the fitted function gives *k* = 0.110, C_L,0_ = 0.452 and C_D0_ = 0.0270. It is impractical due to constraints on simulation time to perform the same array of CFD simulations for all 42 tail postures, each of which would require simulations at a range of angles of attack. Instead, we simulated a range of angles of attack for just five tail postures (Figure S3) and used these for validation of our drag model.

**Figure 4.**
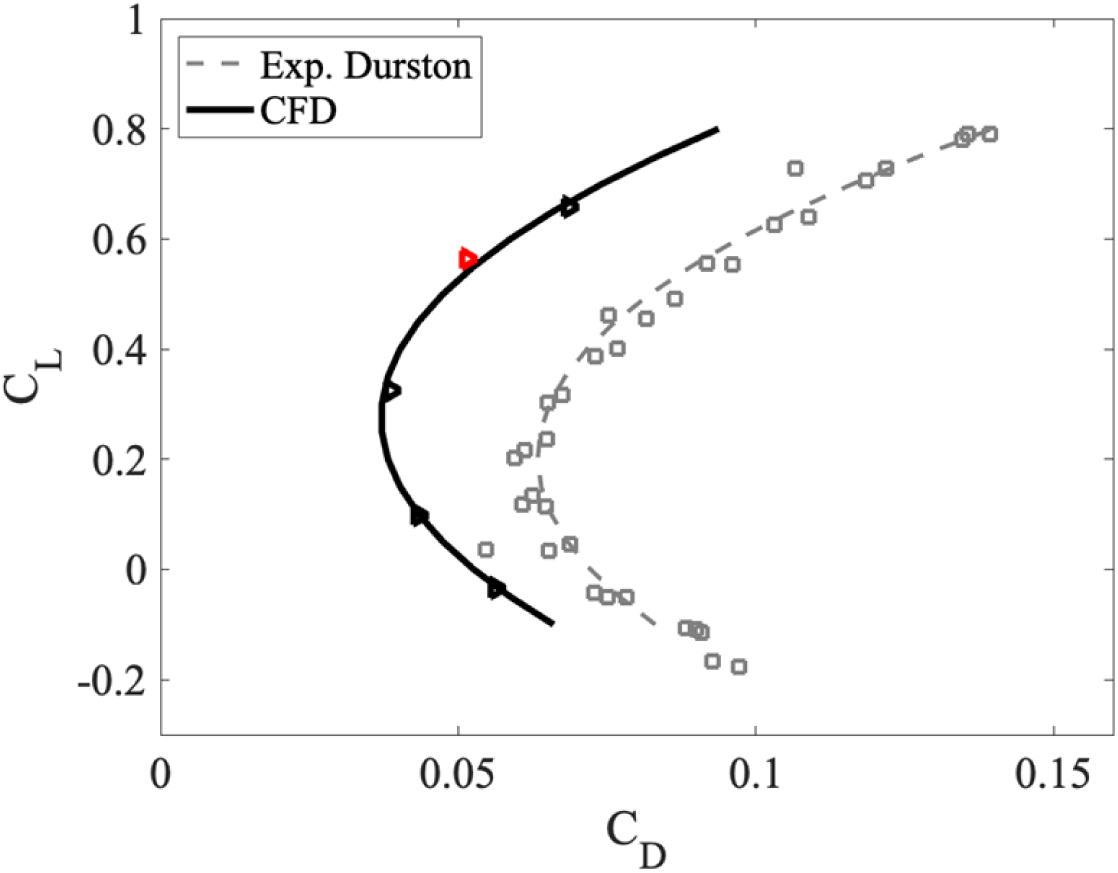
C_L_-C_D_ relation with five angles of attack with 2.5° interval. The red triangle denotes the barn owl with observed posture at its observed angle of attack of 2.9°. The C_L_-C_D_ is fitted with binomial. Wind tunnel measured data from a 3D printed barn owl model is shown for comparison [23], which also shows the offset due to camber.

Since *k* is typically treated as a constant varying only with Reynolds number, its value for different tail postures is maintained at *k* = 0.110 based on the polar from the barn owl’s observed posture. From the C_L_-C_D_ polar in Figure S3, we can see that the symmetric axis of the curve is approximately at C_L_ = 0.2-0.3. Thus, the zero-lift drag can be approximated as the drag directly obtained from CFD simulations. With that, the unknown parameters C_L,0_ and C_D0_ can be derived (Figure S4). The comparison between the C_D0_ using these assumptions and the CFD polar fitting gives, at most, 8.02% difference (Fig. S5 a) and the overall drag differed by 4.39% (Fig. S5 b).

With values of C_L,0_ and C_D0_ estimated using the method above, the drag of all 42 tail postures was determined with the same body weight support (Fig. 5a). Minimum drag now occurs at spread angle 18° and pitch angle 23° when gliding at 7.88 m/s. Of 16 measured glides of the same barn owl, six flights occur at speeds within 2% of 7.88 m/s and are located in the region of minimal total drag. We can delve further into this drag analysis. As described above, viscous drag is the remaining drag after the induced drag has been subtracted from the total drag. Minimal viscous drag is achieved at *ϕ*_*s*_ = 23.2°, *ϕ*_*p*_= 25.6° while minimal induced drag is achieved at *ϕ*_*s*_ = 15.0°, *ϕ*_*p*_= 36.1°. At the measured tail posture, induced drag and viscous drag contribute 32.3% and 67.7% of the total drag, respectively(Fig. 5b and c). Thus, viscous drag is twice as important as inviscid drag for barn owls at these speeds.

**Figure 5.**
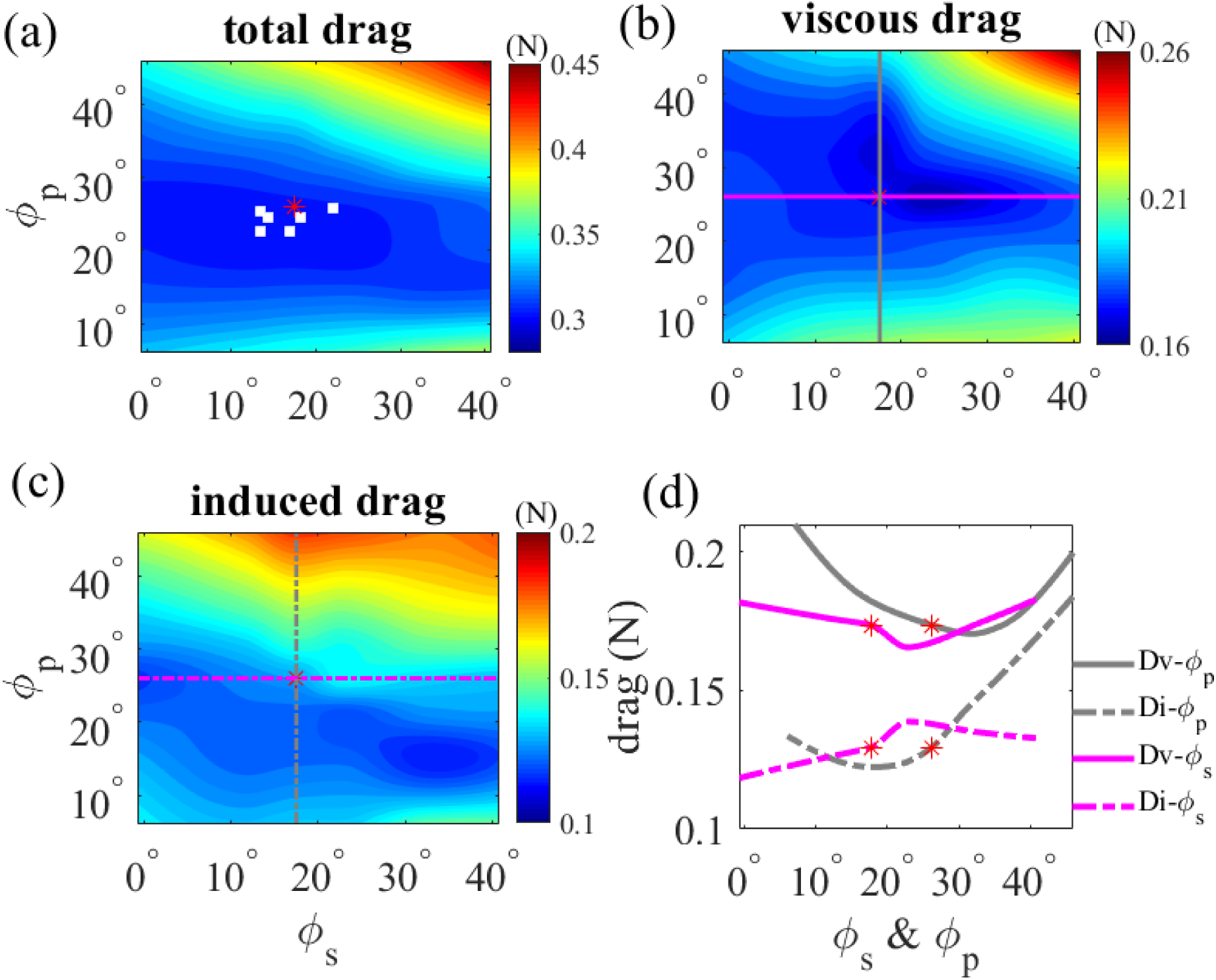
Drag predicted by the drag model with the same body weight support for different tail spread and pitch angles. (a) Total drag, white squares show the tail posture of observed flights within 2% of the speed of the tail posture case used for simulation; (b) Viscous drag; (c) Induced drag. (d) Drag variation as a function of spread and pitch angle originating from the observed posture. Data is extracted in a transects shown in (b) and and coloured accordingly.

Moving away from the tail posture we observed, viscous drag increases more dramatically than the induced drag when the tail pitches up; whilst the induced drag increases more dramatically when the tail pitches down (Fig. 5d). As the minimal viscous drag and induced drag cannot be achieved simultaneously – *i*.*e*., with same tail pitch and spread angles – the barn own appears to be minimising total drag with a compromise tail pitch angle between the minimal viscous drag pitch angle (*ϕ*_*p*_= 35.8°) and the minimal induced drag pitch angle (*ϕ*_*p*_ = 19.8°).

### Variation in drag with flight speed

To give confidence in the robustness of our results and predict changes in tail posture with flight speed, we repeated our drag predictions for a range of tail pitch and spread angles at four additional speeds: 6.5, 7.47, 8.49, and 9.0 m/s. The parameters *k*, e_i_, e_v_ C_*L*,0_ and C_D0_ were treated as independent of flight speed, so C_L_ is the only variable. Generally speaking, at the lowest speed (U = 6.50 m/s) and highest speed (U = 9.00 m/s), drag is larger than the case of U = 7.88 m/s for all postures, indicating that the barn owl chose to glide at an optimal flight speed for minimal drag with that tail posture or, conversely, that tail posture was chosen to minimize drag at that speed (Fig. 6). At each flight speed, there is an alternative tail posture that has lower drag. However, the minimal drag for some speeds (*e*.*g*., U = 6.50, 8.49 and 9.00 m/s) was beyond the range of tail spread and pitch angles in our parameter sweep. Interestingly, within some regions of the tail posture graphs it appears that drag is somewhat insensitive to tail posture. In these regions, the tail could be used for control with little additional cost in terms of flight efficiency.

**Figure 6.**
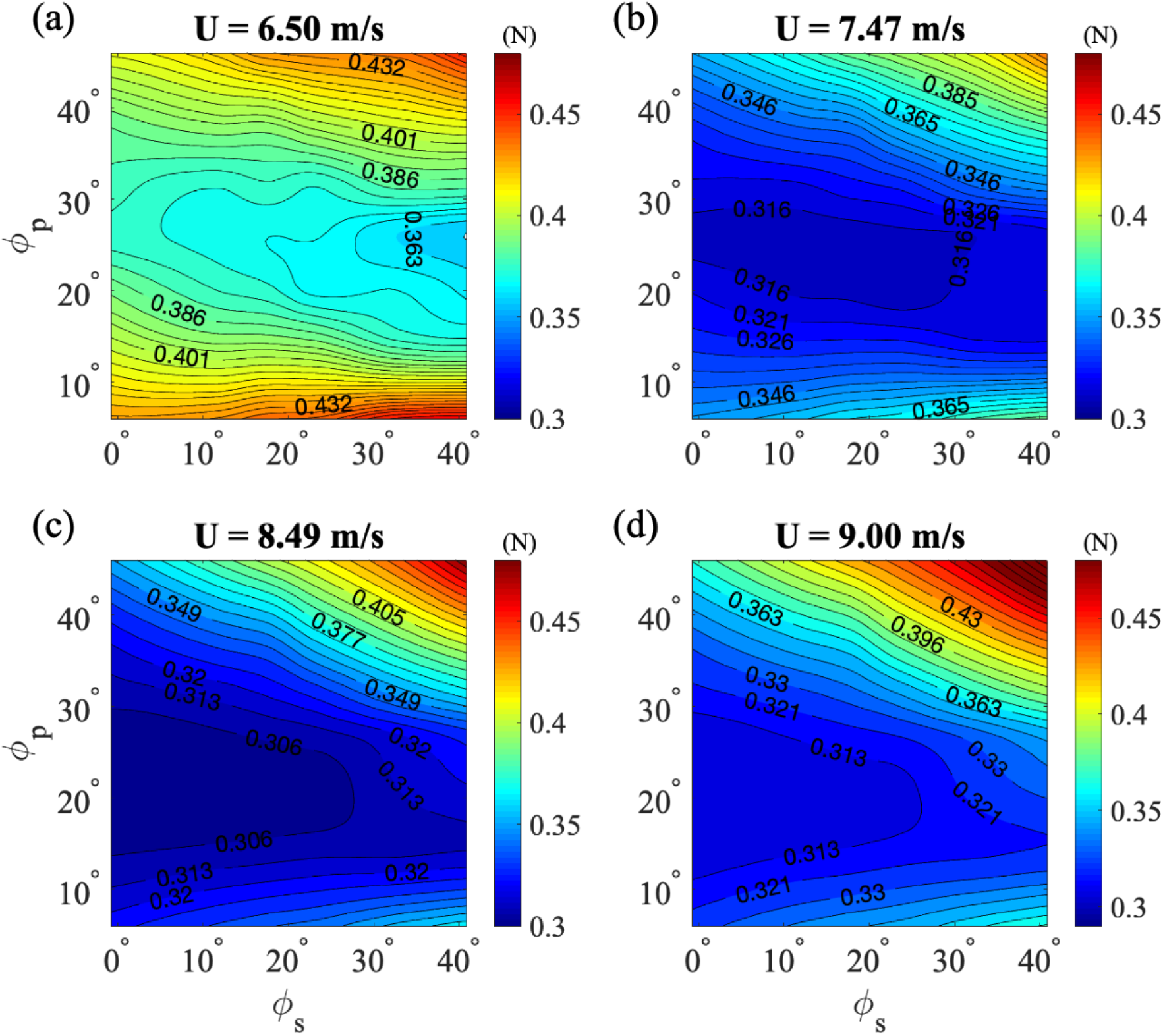
Total drag predicted by the drag model with different tail postures at four flight speeds (U = 6.50m/s, 7.47 m/s 8.49 m/s and 9.00 m/s).

We compared the drag estimated for 16 observed tail postures with the minimal drag prediction from the drag model (Fig. 7). The drag prediction offset b is only 0.85% of the minimal drag, and *κ* = 1.043 shows the drag of the observed tail posture leads to an average of just 4.3% increase over the minimum drag predicted by our model. This supports the notion that the selected tail posture was operating near minimal drag.

**Figure 7.**
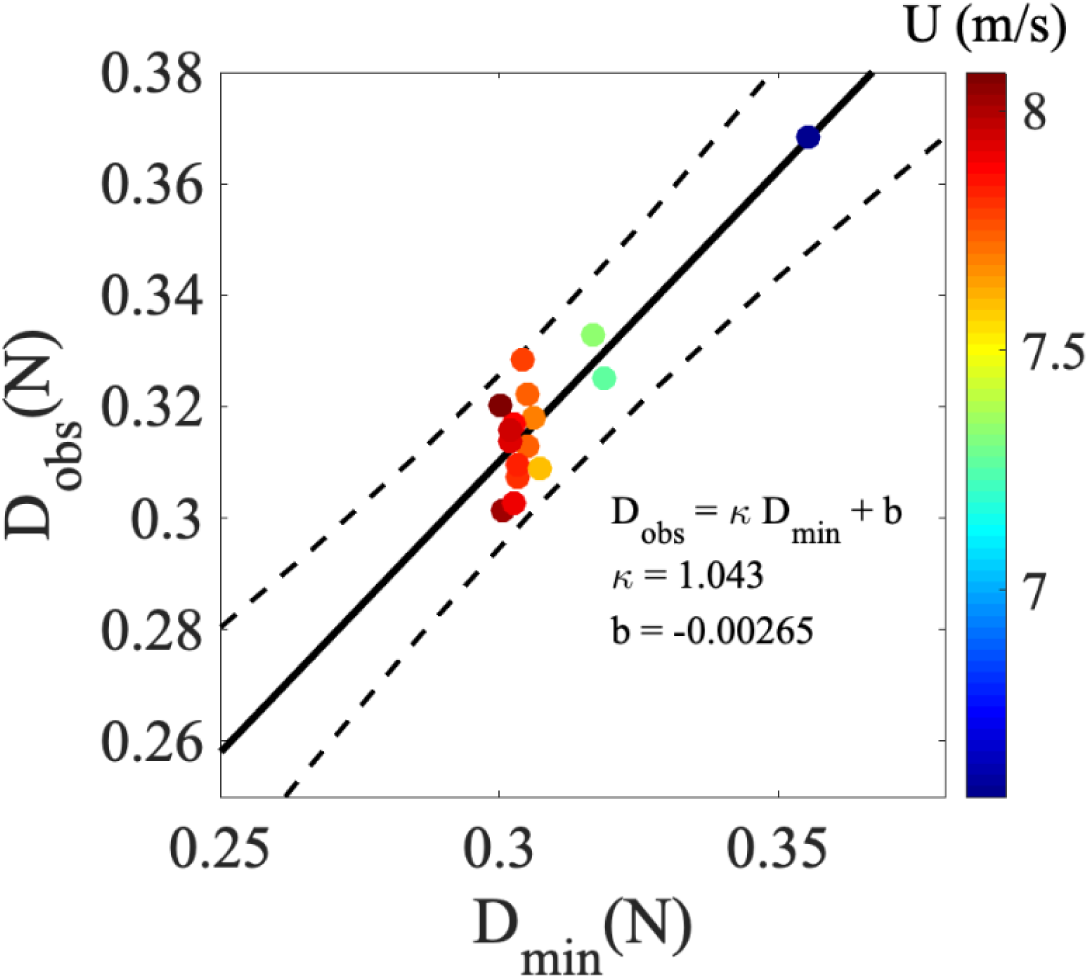
The drag estimated with 16 observed tail postures versus the minimal drag prediction from the drag model. The color of circles denotes the flight speed. The solid line denotes the linear fitted line of observed drag versus theoretical minimal drag using model D_obs_ = *κ*D_min_ + b (*κ* = 1.043, b = −0.00265), and the dashed lines denote the 95% confidence interval.

## Discussion

### Aerodynamic roles of tail

We have described how the tail operates to reduce total drag. Previous studies have shown that at low speed, birds spread their tails widely and operate them at a high angle of attack relative to the flight direction [5,6,29,30] indicating that birds use the tail to generate lift in order to support body weight [10,30]. We found by integrating tail surface pressure that the tail alone contributes just 3% of body weight support, which is a similar value to that measured in a flying starling (2%) [4]. The use of differential pressure sensors on the tails of pigeons during level flight suggested 7.9% weight support [31]. Using stagger theorem and relation for the induced drag of a biplane, Thomas (1996) showed the induced drag of the wing-tail combination is lower than the induced drag of a wing alone provided that the tail generates a positive lift [29]. At a speed of U = 6.50 m/s the barn owl demonstrates low drag with a high spread angle and relatively large pitch down angle. Therefore, the tail can be used to increase lift as well as decrease drag, which means a spread tail at low speed increases the lift/drag ratio.

For a flying bird, a stable configuration might reduce energy consumption during long-distance flight by reducing control inputs [23]. The bird would become more stable longitudinally were the tail to pitch up and provide longitudinal dihedral. However, the negative angle of attack required of the tail would reduce lift and introduce extra drag (Fig. 3b). This posture is not used by the barn owl but can be observed momentarily as pigeons prepare to land [31]. As the tail usually pitches up, a reverse cambered body-tail chord confers longitudinal stability, which suggests the tail can also act as a pitch stabilizer [26].

### Efficiency coefficients

At steady gliding flight, the barn owl has a wash-in wing configuration, with high angle of attack at the wing tips that changes gradually to 0° at the shoulders. The wash-in configuration leads to a large downwash behind the wing tips. The span efficiency coefficient e_i_, a parameter related to downwash evenness, is 0.78 for the observed tail posture case, but could exceed 0.86 if the tail were suitably adjusted in spread and pitch. In comparison, the gliding swift gives e_i_ = 0.56 [30], which is lower than our barn owl; the estimation of three raptors using constant-Cl theoretical downwash profiles gives e_i_ 0.8∼0.9 [10].

Viscous efficiency defines the proportion of drag that, due to poor loading distribution, exceeds the minimum ‘useful’ drag, *i*.*e*., the minimum viscous drag associated with successful overall weight support of a wing with each section at ideal (and potentially different) incidence. A full calculation of this requires an involved optimisation that is beyond the scope of this study; instead, we assume sectional aerofoil properties are constant. In this case, the highest viscous efficiency (e_v_ = 1) can be achieved only if the lift coefficient of 2-dimensional chords across the span is constant. The spanwise distribution of chord length in birds varies considerably (as the torso/tail region is much longer than the wing cross sections) and, as a result, the 2-dimensional force at the torso/tail region should be much larger to maximise viscous efficiency. This leads to a strong downward jet behind the torso/tail [10]. Throughout the range of the tail manipulation, e_v_ ranges from e_v_ = 0.77, when the tail is pitched-up, to e_v_ = 0.99, when the tail is spread and pitched-down. There is a paucity of data in the literature suitable for calculating e_v_, but the strong downwash behind the tail evident from flow visualisations implies a relatively high e_v_ [10,30], which is consistent with the value e_v_ = 0.88 for our observed tail posture. However, note that, as we adopt the 2-dimensional flow assumption, so the analysis in this paper is less applicable to birds with particularly long necks or extremely long tails.

As high e_i_ is achieved when the lift distribution across the span is elliptical, and high e_v_ is achieved when the lift coefficient is constant (*e*.*g*., lift distribution is proportional to wing chord), e_i_ and e_v_ can’t be optimised simultaneously for non-elliptical avian planforms. The e_i_-e_v_ relation for a simplified wing planform can be estimated using lifting-line theory [32] (Fig. 8). For this notional non-elliptical planform, e_i_ and e_v_ are in a competitive relationship dependent on the lift distribution. Dominance of one efficiency factor over another can be prescribed by manipulating the tail’s pitch and spread angles. The overall aerodynamic performance depends on the relative significance of the induced and viscous drag contributions to the total drag on the bird; high e_i_ should be favoured when induced drag dominates at high speeds and/or large scales, while high e_v_ should be favoured in circumstances of low speeds and/or small scales where viscous drag dominates.

**Figure 8.**
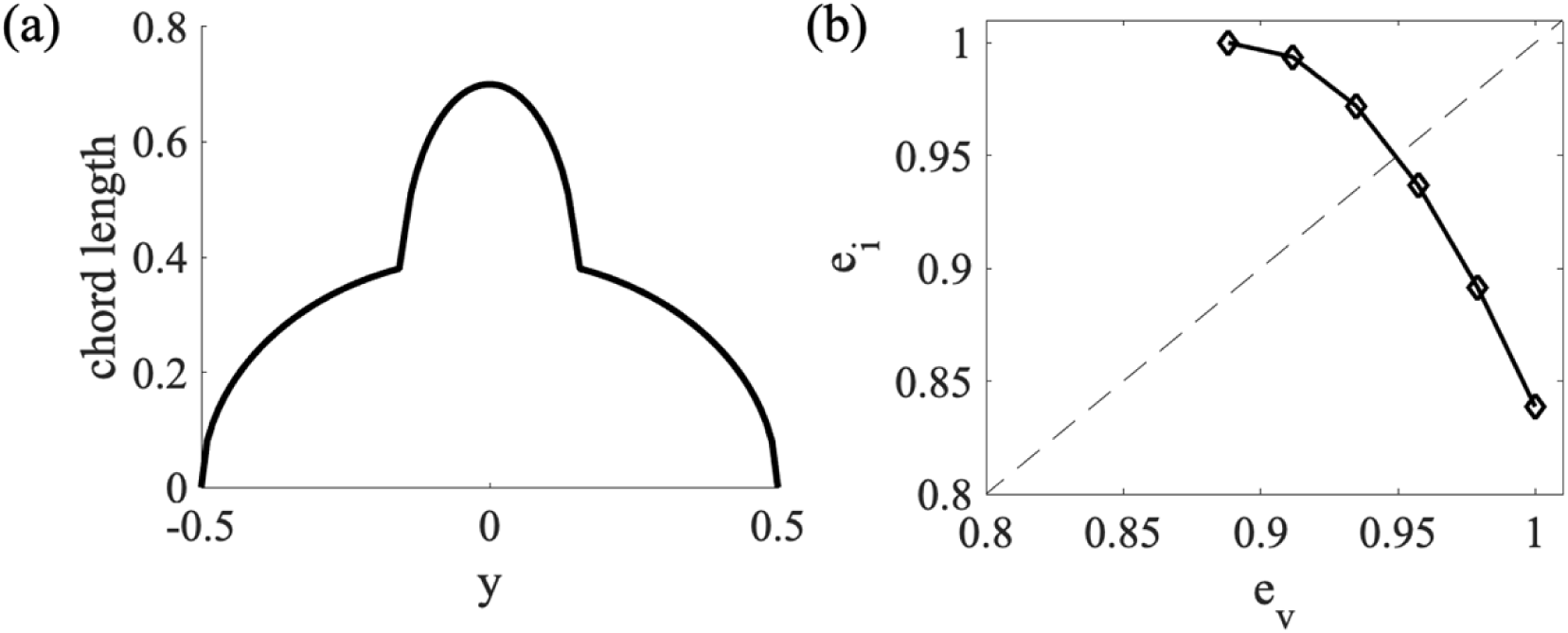
The relationship of e_i_ and e_v_ for a notional planform of a bird with a tail. The left figure shows the chord distribution of the planform. The right figure shows how e_i_ and e_v_ vary due to the manipulation of tail’s pitch and spread angles.

### Variation with speed

Wind tunnel studies have shown that the wing span of gliding animals decreases as the flight speed increases [5,14,15]. An investigation of a jackdaw [15] showed no noticeable effect of spanwise camber on e_i_ due to speed variation. Substituting C_L_ = L/*qS* ≈ W/*qS* into Eq. 8 gives,

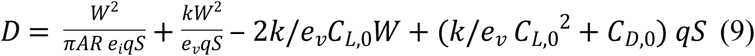

and we can see that the overall drag will be minimal at a certain flight velocity (*q* = 0.5 *ρU*^*2*^). With this drag model, we can easily estimate the gliding polar of the bird (Fig. 9). This polar reveals the speed giving minimum sink rate to be, U_ms_ = 7.74 m/s, while the speed for maximum range is 12% faster at, U_bg_ = 8.68 m/s. As 9 out of 16 observed flights were within 2% of U_ms_, we can conclude that the barn owl chose a speed that would minimize the sinking rate when gliding through the flight corridor.

**Figure 9.**
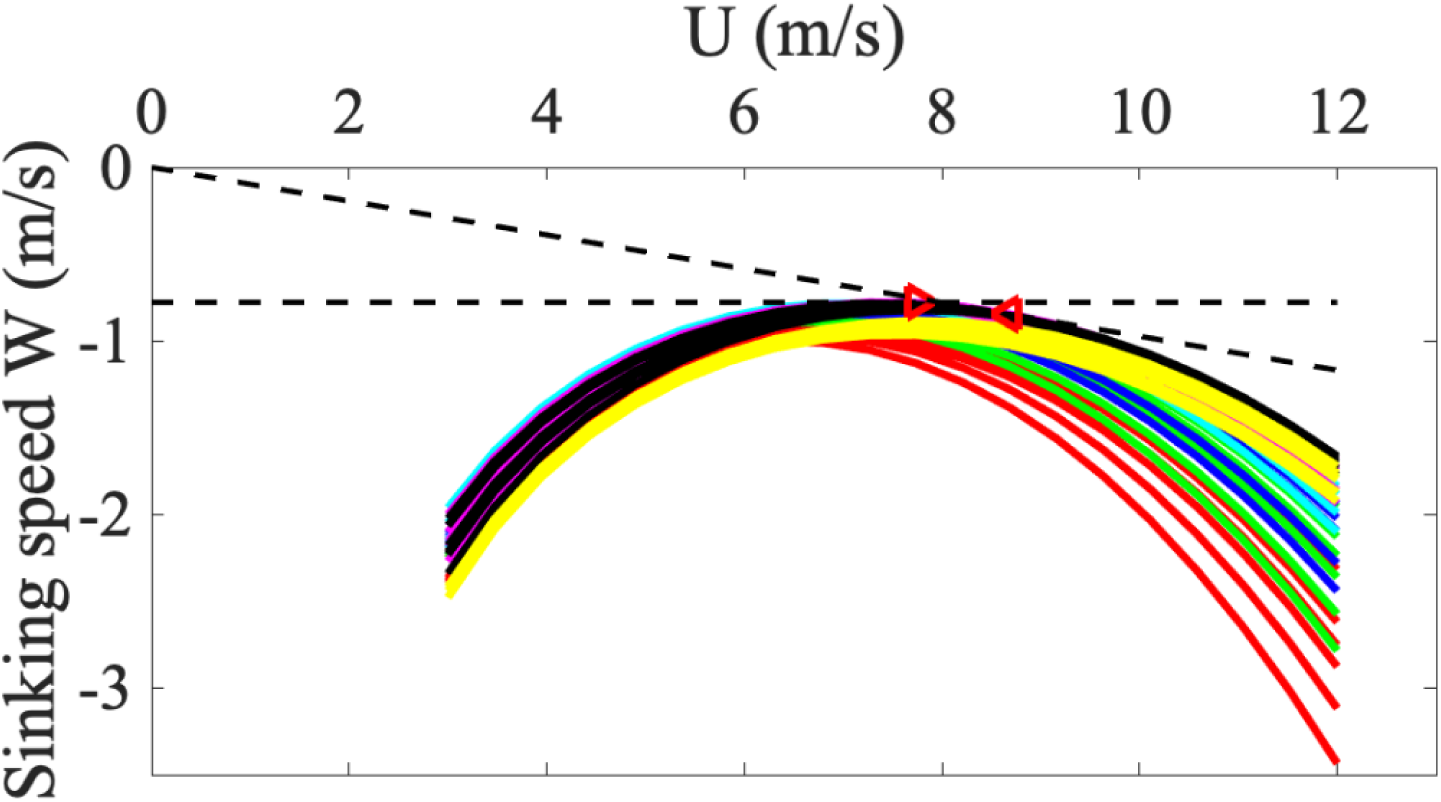
gliding polar of a barn owl. Lines denote the polar for different tail postures, the lines with same color denote the case with same tail pitch. Right oriented triangle ((U, W) = (7.74, 0.778) m/s) denotes the speed giving minimal sinking rate while left oriented triangle ((U, W) = (8.68, 0.846) m/s) denotes the speed giving maximum gliding range.

**Figure 10.**
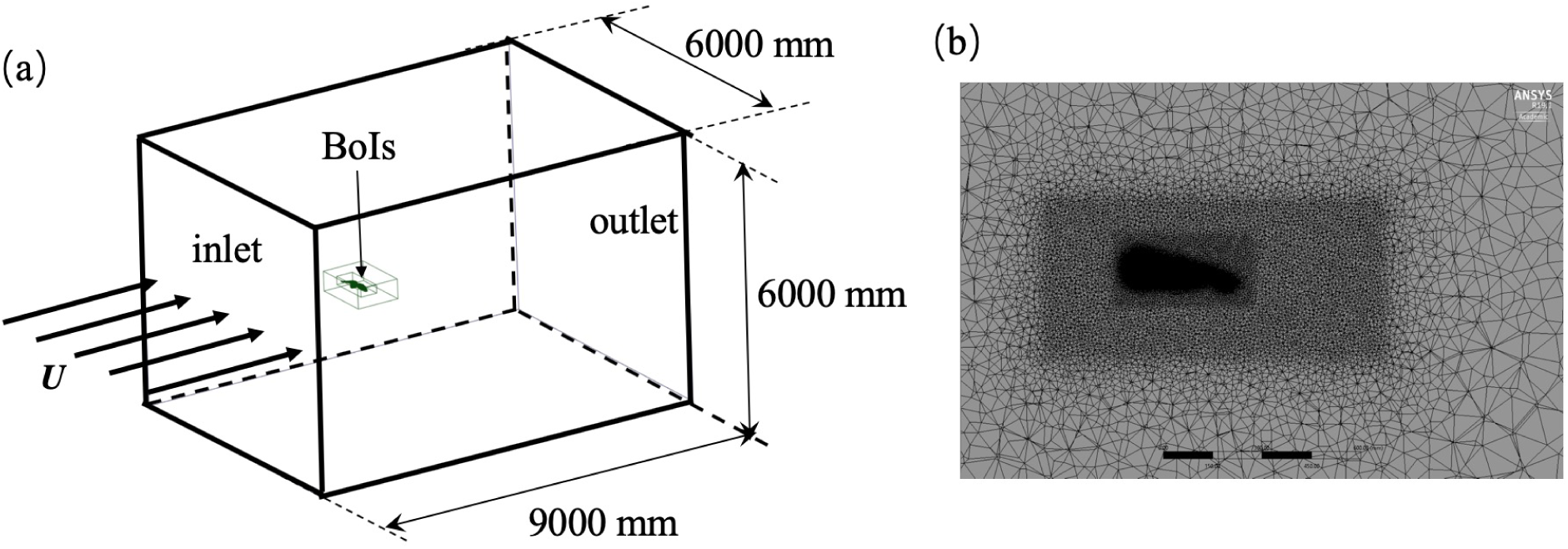
CFD simulation setup (a) and the mesh on the sagittal plane of the computational domain (b). Each black scale bar in (b) denotes the length of 150 mm.

### The effect of *k*

In the 2-dimensional drag model, the parameter *k* describes the quadratic rise in C_d_ with C_l_ – C_l,0,_ and varies with Reynolds number. It has been suggested that the same *k* can be used for both 2-dimensional and 3-dimensional wings [25], which means the *k* for any chord is constant. During the derivation of the comprehensive 3-dimensional drag model, we adopted this assumption. The determination of *k* is based on fitting the 3-dimensional drag model to the CFD simulated C_L_-C_D_ polar. Aside from obtaining *k* from the observed tail case (*k* = 0.110), the other four postures gave values of *k* = 0.0845, 0.0839, 0.0903 and 0.114. Using the e_i_ and e_v_ values from the observed tail posture case, fitting the model to C_L_-C_D_ polar measured from a 3D-printed barn owl model gives *k* = 0.114 [23], similar to our result. Across birds, variation in *k* is relatively small: using data for 20 birds from previously published papers [33,34], we estimate the *k* value to be 0.0926 at similar Reynolds numbers to our barn owl. Therefore, across birds we can see that variation in *k* between species is not large. In addition, the total drag is relatively insensitive to variation of *k* in the model, when lift equals weight support. The change in drag coefficient with respect to change of *k* can be calculated as follows:

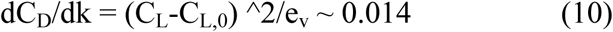

or, alternatively, we find a 0.03% change in drag coefficient for every percent change in *k*. The dominant effects of the minimum drag and inviscid drag for our reference glide make the model insensitive to *k*.

## Conclusions

In this paper, we developed a comprehensive analytical drag model, calibrated by high-fidelity computational fluid dynamics (CFD), to investigate the aerodynamic action of the tail by virtually manipulating its posture. Using this CFD-calibrated drag model, we predicted changes in drag when varying the tail pitch and spread angles while maintaining body weight support. By comparing the minimum predicted drag that was possible within a wide range of tail postures against the drag calculated for 16 observed gliding flights, we found that the observed postures of our gliding bird corresponded to near minimal total drag. This suggests the bird adjusts its tail posture for minimal total drag and the mechanism by which it achieves that performance is by compromising between the ideal postures required to minimise the induced and viscous components of the total drag. This result shows the mechanism by which birds’ tails, that are most conspicuously used for manoeuvring or trim, can also be used to reduce the burden of drag. It also has significance for improving the flight efficiency of bird-size air vehicles.

## Materials and Methods

### Geometry from stereo-photogrammetry and mesh reconstruction

The solid-body geometry of the barn owl in free flight was obtained from Cheney and colleagues [12], and this approach is also utilized by Durston and colleagues [26]. The solid body was reconstructed from an indoor glide of a barn owl using photogrammetry and twelve high-speed cameras. The same fluid mesh was used as in this study, this fluid mesh shows good agreement when comparing the output against weight support and downwash profile. This work was approved by the Ethics and Welfare Committee of the Royal Veterinary College (URN2018 1836-3).

### CFD simulation

The simulation domain was 9000 × 6000 × 6000 mm3, and the bird model was placed 3000 mm downstream from the inlet (Fig. 9a). The fluid domain mesh was generated by ANSYS Mesh 19.1 (ANSYS, Inc., Canonsburg, PA, USA). We used a hybrid mesh, including tetrahedral, pentahedral and hexahedral cells with multiple body of influences (BoIs) to discretize the fluid domain (Fig. 9b). Adjacent to the bird surface, the inflation layer had a first layer thickness of *δ*_*t*_ = 0.1 mm (y+=3), a growth factor of 1.2, and 19 layers. Mesh independence is shown in Table. 3, justifying the use of *δ*_*t*_ = 0.1 mm in this study. Two bodies of influence were used to control mesh size, with the mesh size in the inner domain being 5 mm and the outer domain being 12 mm.

Commercial software FLUENT 19.1 (ANSYS, Inc., Canonsburg, PA, USA) solved the fluid dynamics governing equations using a two equation *k* − *ω* SST turbulence model. The velocity at inlet was constrained to 7.88 m/s (U) (Fig. 9a), and at outlet followed the Neumann boundary condition. Pressure at inlet and outlet followed the Neumann boundary condition as well. The far-field boundary conditions, perpendicular to the flow, were constrained to be symmetric. Turbulence intensity was specified as 1% at the inlet to simulate the turbulence of flow in the flight corridor, which we expect is greater than in a high-quality wind tunnel.

### Solving for aerodynamic model coefficients using CFD

We computed 62 fluid simulations around the bird model: 42 simulations were at the same angle of attack with systematically varying tail pitch and spread, the additional 20 simulations were to compute the aerodynamic polar for the reference tail-posture case, and four additional tail postures bounding the observed flight data (Figure S4).

The five unknown aerodynamic coefficients in our model are the viscous and inviscid efficiency (e_v_ and e_i_), empirically-derived curvature of the polar (*k*), minimum-drag coefficient (C_D0_), and camber-induced-lift coefficient (C_L,0_). For all flights, we solve for viscous and inviscid efficiency using the distributions of lift coefficients and downwash, respectively (Eq. 6 and 10). We assume polar curvature is negligibly different across tail morphology, and use the quantity derived from the reference tail-posture case, which is roughly centred in our pitch-spread parameter sweep. Tor the five cases with additional aerodynamic polar data, we solved for the minimum-drag coefficient and camber-induced-lift coefficient by fitting using the linear, least-squares method. We observed that for each of these five cases, drag produced at the natural glide angle was roughly equal to the computed drag at zero lift; we do not believe that this has to be a general phenomenon, but use it here as a convenient model heuristic. This allows us to compute camber-induced lift and minimum drag, for flights with a single point on the aerodynamic polar, by adding the assumption that drag at zero lift equals drag at the natural glide angle.

### Span efficiency e_i_

Span efficiency is calculated by the equation

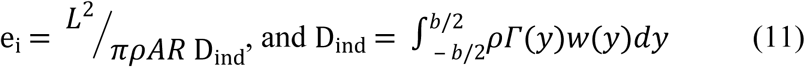

where *Γ*(*y*) and *w*(*y*) is the wing boundary circulation and downwash distribution along the span, respectively [35].

## Acknowledgements

We thank Masateru Maeda for helpful discussions. J. S. thanks Profs. Hongwei Ma and Zhenzhong Sun for providing computing resources in DGUT.

## Supporting information

**S1 Figure**. Histogram of the distance between the point clouds and the surface mesh reconstructed from the point clouds.

**S2 Figure**. Comparison of turbulence models. All models give similar results for lift. In comparison, there is greater variation in the estimated drag, especially for the two *k* − *ε* models, which are known to perform poorly in curved boundary layer simulations.

**S3 Figure**. (a) CL-CD plot of five tail postures, comparing to the data measured on a model barn owl. The five postures are shown in (c). (b) shows the enlarged plot selected from (a). Yellow filled circles denote the C_L_ and C_D_ where the body weight is supported.

**S4 Figure**. C_L,0_ and C_D0_ variation as the function of tail spread and pitch angle.

**S5 Figure**. (a) Comparison of the CD0 calculated from the CFD polar plot and the one used in drag model. (b) Total drag calculated from CFD and estimated from the calibrated drag model.

**S1 Video**. Error distribution of the reconstructed barn owl geometry. The error bar ranges from - 4.5 mm to 4.5 mm. 96% of the points are within the 3.0 mm error, and 90% of the points are within the 2.0 mm error.

**S2 Video**. Virtually manipulated tail animation. The tail spread and pitch are actively changed to certain values using a linear mapping by MATLAB. Subfigures show the detailed changes in three orthogonal perspectives.

**S3 Video**. Variation of pressure deviation from the observed tail posture case on the bird surface due to the tail posture manipulation, *e*.*g*., the subtraction of the pressure on the bird surface by the pressure distribution of the bird with observed tail posture.

